# Large-scale deregulation of gene expression by artificial light at night in tadpoles of common toads

**DOI:** 10.1101/2021.07.08.451570

**Authors:** Morgane Touzot, Tristan Lefebure, Thierry Lengagne, Jean Secondi, Adeline Dumet, Lara Konecny-Dupre, Philippe Veber, Vincent Navratil, Claude Duchamp, Nathalie Mondy

**Affiliations:** Univ Lyon, Université Claude Bernard Lyon 1, CNRS, ENTPE, UMR 5023 LEHNA, F-69622, Villeurbanne, France; Faculté des Sciences, Université d’Angers, 49045 Angers.; Université de Lyon, Université Lyon 1, CNRS, Laboratoire de Biométrie et Biologie Evolutive UMR 5558, F-69622, Villeurbanne, France; PRABI, Rhône-Alpes Bioinformatics Center, Université Lyon 1, 69622 Villeurbanne, France; Institut Français de Bioinformatique, UMS 3601, 91057 Évry, France

## Abstract

Artificial light at night (ALAN) affects numerous physiological and behavioural mechanisms in various species by potentially disturbing circadian timekeeping systems. Although gene-specific approaches have already shown the deleterious effect of ALAN on the circadian clock, immunity and reproduction, large-scale transcriptomic approaches with ecologically relevant light levels are still lacking to assess the global impact of ALAN on biological processes. Moreover, studies have focused mainly on variations in gene expression during the night in the presence of ALAN but never during the day. In a controlled laboratory experiment, transcriptome sequencing of *Bufo bufo* tadpoles revealed that ALAN affected gene expression at both night and daytime with a dose-dependent effect and globally induced a downregulation of genes. ALAN effects were detected at very low levels of illuminance (0.1 lux) and affected mainly genes related to the innate immune system and, to a lesser extend to lipid metabolism. These results indicate that a broad range of physiological pathways is impacted at the molecular level by very low levels of ALAN potentially resulting in reduced survival under environmental immune challenges.

## Introduction

Photoperiod is one of the major environmental cues that regulates biological processes in living organisms (Longcore & Rich, 2004; Ouyang et al., 2017) and allows them to anticipate daily and seasonal changes and adapt their biochemical, physiological and behavioural activities. In the last century, artificial light at night (ALAN), generated by the surge in urbanized areas and transport infrastructures (Gaston et al., 2013), has dramatically expanded worldwide (Falchi et al., 2016) and endangered biodiversity on the global scale (Gaston et al., 2015; Guetté et al., 2018; Hölker et al., 2010; Secondi et al., 2019).

In the last decade, numerous studies have reported the effects of ALAN on many physiological processes in a large panel of animals. This anthropogenic light modifies illuminance levels so that the perception of fine changes in natural light level can be impaired, thus desynchronizing daily or seasonal activities (de Jong et al., 2016; Robert et al., 2015). Hence, the internal circadian timekeeping system is impacted by ALAN mainly through alterations in the expression of clock genes (Bedrosian et al., 2013; Fonken et al., 2013; Honnen et al., 2016, 2019). However, many other genes that are regulated by photoperiod, such as genes related to metabolic detoxification, immunity and nutrient sensing, could also be impacted by ALAN (Honnen et al., 2016). In many taxa, ALAN reduces the nocturnal expression of melatonin (Grubisic et al., 2019), a hormone involved in the regulation of the immune system, energy metabolism, antioxidant defences and sex hormone synthesis (Reiter et al., 2010; Zhao et al., 2019). Short-term exposure to ALAN was, however, not associated with alterations in immunity or oxidative stress in adult fishes (Kupprat et al., 2021) and birds (Raap, Casasole, Costantini, et al., 2016), suggesting that longer exposure and/or a more vulnerable developmental stage might be required to have deleterious effects if any. Accordingly, in humans and rodents, prolonged ALAN exposure is associated with an increased prevalence of obesity, type 2 diabetes, hypertension, affective disorders or specific cancers (Fonken & Nelson, 2014; Haim & Portnov, 2013; Wyse et al., 2011).

At the molecular level, the impact of ALAN is beginning to be studied, but the studies target mainly specific candidate genes (Bedrosian et al., 2013; Brüning et al., 2018; Fonken et al., 2013; Honnen et al., 2019). However, given the multiplicity of ALAN effects on organisms, integrative approaches are required to simultaneously investigate all these effects at the gene level and better understand the causes of physiological and behavioural changes triggered by this stressor. A pioneering study using a large-scale transcriptomic approach revealed a sex-specific response of urban adult mosquitoes to ALAN, with reduced expression in males of genes mostly related to gametogenesis, lipid metabolism and immunity (Honnen et al., 2016). A second study characterizing the transcriptomes of coral reefs showed alterations of the pathways associated with the cell cycle, growth, proliferation and protein synthesis in ALAN-exposed coral reefs (Rosenberg et al., 2019). However, both studies used rather high experimental light levels to mimic direct illumination at night. To our knowledge, no transcriptomic study has yet been conducted using low light levels representative of levels recorded in peri-urban or rural areas, although the low light levels would be more relevant from an ecological point of view (Luginbuhl et al., 2014). In addition, while transcriptomic analysis of ALAN impacts was restricted to the night period, several studies showed that ALAN had long-lasting impacts on animal physiology and behaviour during the daytime (Dominoni et al., 2020; Latchem et al., 2021).

Freshwater ecosystems are particularly concerned with ALAN exposure (Grubisic, 2018; Secondi et al., 2017) since lit road networks, urban development and industrial infrastructure are frequently located along rivers and lakes (Reid et al., 2019). However, these habitats are home to rich fauna (Dudgeon et al., 2006), which will thus suffer the effects of ALAN. This is particularly the case for amphibians, such as the common toad, *Bufo bufo.* This species is one of the most common amphibians in western Europe, and frequently colonizes urban areas (Beebee, 1979). In the present study, we assessed transcriptome-wide gene expression in response to ALAN in tadpoles of the common toad. We combined *de novo* transcriptome sequencing and assembly and a controlled laboratory experiment to investigate the transcriptome-wide gene expression response using Illumina RNA-seq in common toad tadpoles following prolonged exposure to ALAN. We used three ecologically relevant ALAN levels at night: 0.01 lux (control), 0.1 lux, which corresponds to the illuminance of urban skyglow (Gaston et al., 2013), or 5 lux, which corresponds to the light level of a residential street (Gaston et al., 2013). After 27 days of exposure, the ALAN effect was evaluated for each light treatment both at night (01:00) and daytime (13:00).

## Results

### ALAN affected gene expression in *B. bufo* tadpoles

The sequencing of 30 individual tadpoles bred experimentally with nocturnal light mimicking three levels of ALAN (0.01, 0.1 or 5 lux) and randomly sampled during night-time or daytime yielded a total of 863.8 million 50 bp single-end reads, averaging approximately 28.8 million reads per sample (range 19,700,000 – 38,700,000 reads). Since no genetic data were available for this non-model species, we first assembled a *de novo* transcriptome from all sequenced RNA from *B. bufo* samples, which yielded 38,692 contigs (coding sequences *i.e.,* genes) and 95,483 total transcripts (isoforms).

A principal component analysis (PCA) representation of the transcriptome-wide gene expression variations clearly discriminated the samples according to nocturnal illuminance treatment and sampling period (Fig 1). This global representation showed that control individuals are separated from the individuals exposed to 5 lux, indicating a clear effect of ALAN on gene expression. Moreover, within these two light illuminances, a day/night effect was observable, distinctly separating individuals collected during night-time from those collected during daytime (Fig 1). The samples from the 0.1-lux group overlapped both with the 5-lux group at night and with the control group in the daytime (Fig 1). This complex distribution indicated a different effect of 0.1 lux on day/night gene expression, suggesting that at a light intensity of 0.1 lux, major effects occurred at night, whereas with a higher light intensity of 5 lux, gene expression was affected regardless of the time period.

**Fig 1:**
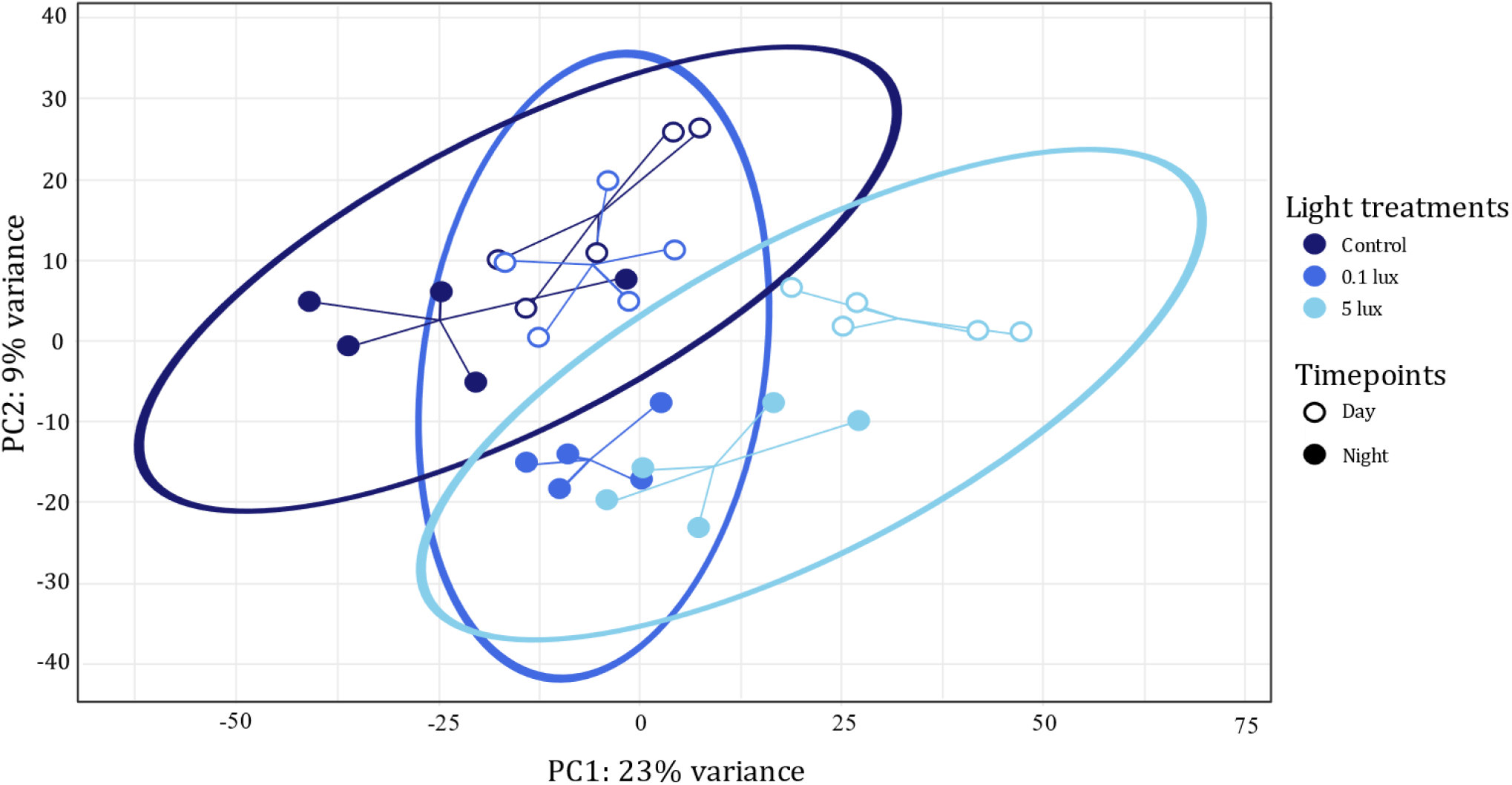
PCA plot of the 30 *B. bufo* transcriptomes. Individuals were exposed to one of three different light treatments: control (dark blue), 0.1 lux (royal blue) or 5 lux (light blue) and sampled at different timepoints: daytime (open symbols) or night-time (closed symbols). Each point represents one individual sample and ellipses represents a 95% confidence level for the multivariate normal distribution of samples in a light treatment. Distance of points to its centroid according to light treatment and day/night effect is represented by a line. PCA plot was illustrated on the two principal axes, explaining 32% of the variability.

### ALAN altered nocturnal gene expression regardless of the illuminance level

When focusing on the variation in gene expression at night, we showed that exposure to ALAN at both 0.1 and 5 lux induced differential expression of genes compared to controls (Fig 2A). The higher the ALAN illuminance, the greater the number of differentially expressed genes compared to controls. Indeed, 3.5% (1,194 genes) and 11% (3,676 genes) of the tested genes were differentially expressed (false discovery rate, FDR < 0.05) in the 0.1- and 5-lux groups compared to the controls, respectively (Fig 2Aa). Among those genes, 284 and 786 had an absolute log2 fold change (|LFC|) greater than 1, and 79 and 82 had an |LFC| greater than 2 in the 0.1- and 5-lux groups, respectively. However, and in accordance with the PCA (Fig 1), far fewer genes were differentially expressed at night between the two ALAN treatments (331 genes with an FDR < 0.05, 125 of which had an |LFC| greater than 1 and 51 of which had an |LFC| greater than 2) (Fig 2Aa). Furthermore, among the 1,194 genes affected by the 0.1-lux group, 76% were also affected by the 5-lux exposure. Among the genes differentially expressed between light treatments at night compared to controls, the majority were downregulated (58% and 62% in the 0.1- and 5-lux groups when compared to controls, respectively) (Fig 2Ab). Once again, the higher the illuminance at night, the greater the under- and overexpression of differentially expressed genes at night compared to controls.

**Fig 2:**
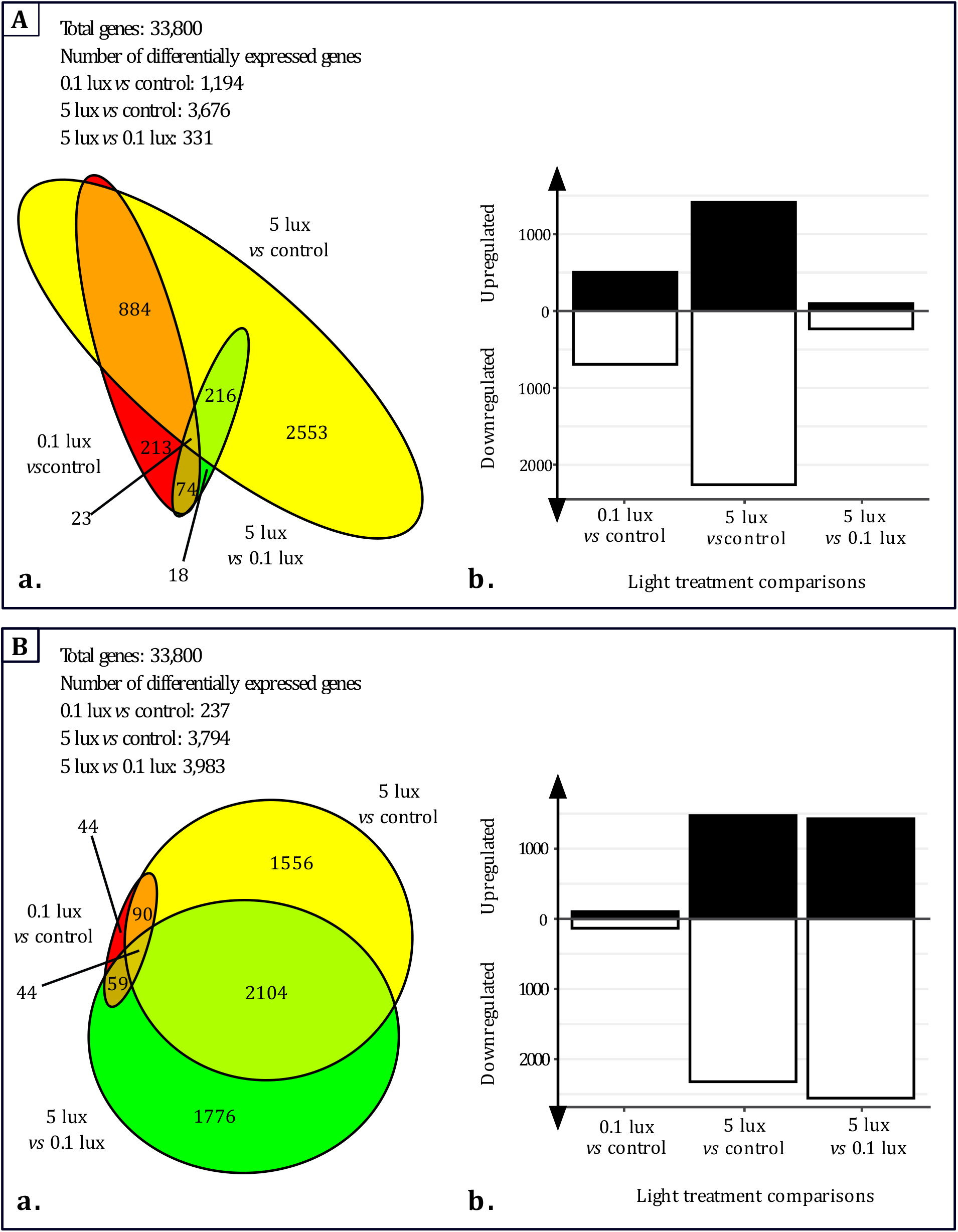
Gene expression variation among light treatments during night-time (A) or daytime (B). a: Venn diagram showing the number of differentially expressed genes (FDR < 0.05) for the different light treatment comparisons. b: Bar plot representing the number of up- (closed bar plot) or downregulated (open bar plot) genes (FDR < 0.05) for the different light treatment comparisons.

### ALAN also altered diurnal gene expression

Our results showed that exposure to 5 lux at night resulted in differential gene expression during the daytime relative to the controls and 0.1-lux group (Fig 2B). We observed differential expression in 11.2% (3,794 genes) and 11.8% (3,983 genes) of the tested genes (FDR < 0.05) in individuals exposed to 5 lux relative to controls or 0.1 lux, respectively (Fig 2Ba). Among those genes, 808 and 952 had an |LFC| greater than 1, and 117 and 153 had an |LFC| greater than 2 in tadpoles exposed to 5 lux compared to controls or 0.1 lux, respectively. Moreover, among the 3,794 affected genes when comparing the 5-lux group to controls, 55.5% were also differentially expressed between 5 lux and 0.1 lux. However, and in accordance with the PCA (Fig 1), far fewer genes were differentially expressed during daytime when comparing the 0.1-lux group to the controls (237 genes with an FDR < 0.05, 90 of which had an |LFC| greater than 1 and 42 of which had an |LFC| greater than 2) (Fig 2Ba). Furthermore, among the differentially expressed genes during the daytime between light treatments, we again observed an excess of downregulated genes in the 5-lux group (61.2% and 64.2% were downregulated when compared to the controls or the 0.1-lux group, respectively) (Fig 2Bb).

### ALAN exposure affected different physiological pathways, mainly the immune system

Functional annotations and overrepresentation of Gene Ontology (GO) terms were analysed on the most strongly differentially expressed genes (FDR < 0.05 and |LFC| > 1) of the 30 samples. The analysis of the GO term enrichment showed that several physiological pathways were affected by ALAN and revealed that genes linked to the immune system corresponded to the physiological pathway mainly affected by ALAN (Fig 3 and S1 Table). Indeed, among the 54 most enriched GO terms (FDR < 0.01 and odds-ratio > 10 for at least one light treatment comparison), 17 were linked to the immune system (Fig 3 and S1 Table). The overrepresentation of GO terms linked to the immune system was observable at night for all light treatment comparisons, even when comparing the 5-lux to the 0.1-lux group (Fig 3 and S1 Table). Indeed, at night, enriched GO terms included antibacterial humoral response and humoral immune response when comparing the 0.1-lux group to the controls. When we compared the 5-lux group to the controls or the 0.1-lux group, the 17 GO terms linked to the immune system were enriched, except for the GO term antibacterial humoral response when comparing the 5-lux group to the 0.1-lux group. The same result was observable during the daytime, with the 17 GO terms related to the immune system that were enriched when comparing the 5-lux group to the controls or the 0.1 lux group (Fig 3 and S1 Table). Globally, at night or daytime, the genes involved in immunity were always downregulated under the higher light level, regardless of the light treatment comparison (0.1 lux *vs* controls, 5 lux *vs* controls or 5 lux *vs* 0.1 lux) (S2 Table). More specifically, the genes that were downregulated at night under ALAN were, for instance, coding for complement C3, complement C3 alpha chain and complement factor H, therefore indicating potential impairment of the innate immune system under ALAN.

**Fig 3:**
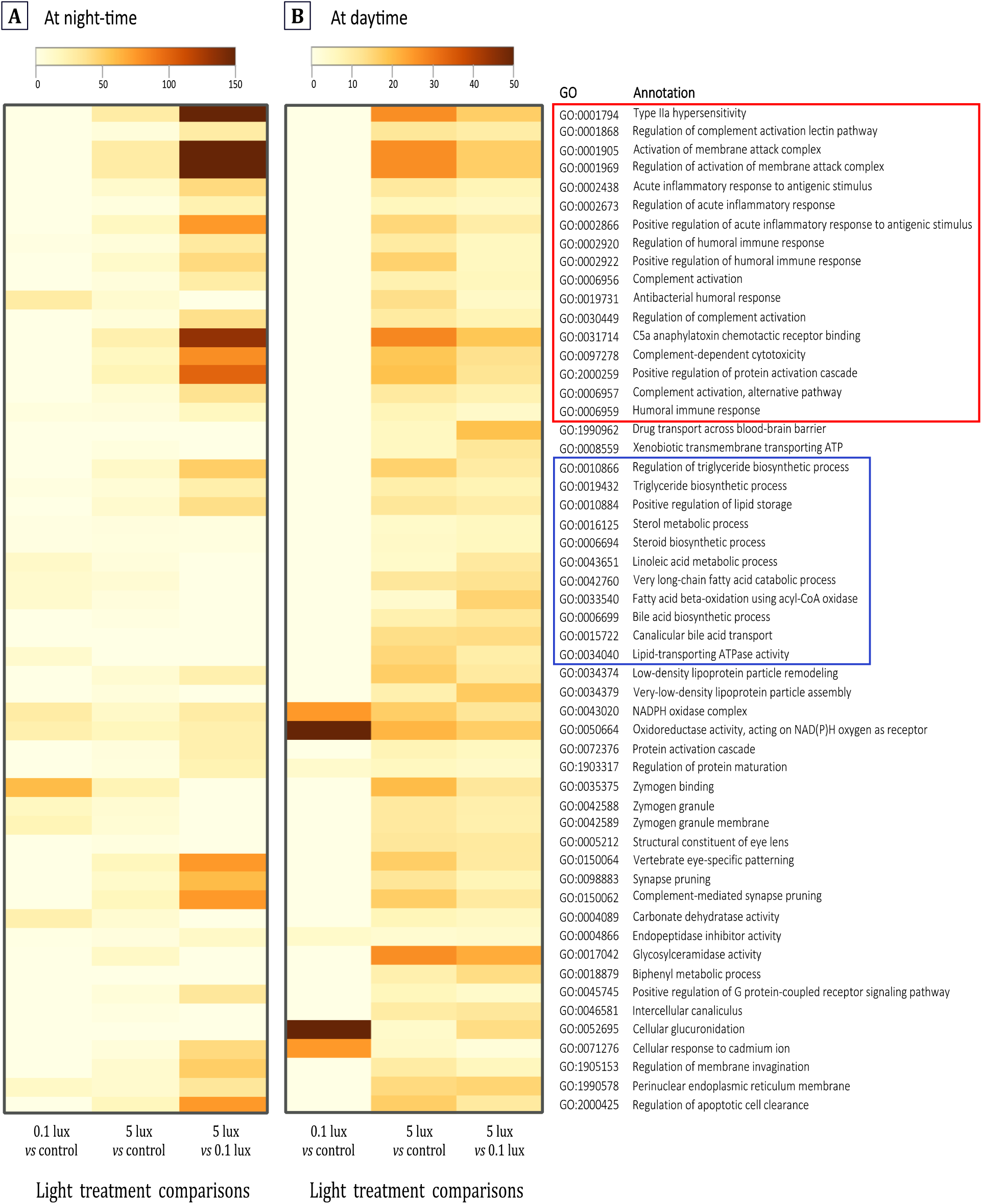
Heatmaps illustrate the enrichment level of the 54 most enriched GO terms (rows, FDR < 0.01 and odds-ratio > 10 for at least one light treatment comparison) for each light treatment comparisons (columns) at either night-time (A) or daytime (B). The colour scale represents the odds-ratio levels; a darker colour indicated a higher enrichment and a lighter colour indicates a lower enrichment. The red insert groups the GO terms associated to the immune system and the blue insert groups the GO terms associated to the lipid metabolism. The data used to generate this heatmap are available in S1 Table.

In addition to the immune system, ALAN also affected lipid metabolism and transport pathways (11 of the 54 most enriched GO terms), involving GO terms such as steroid and triglyceride biosynthetic processes and lipid storage and transport (Fig 3 and S1 Table). Genes involved in these pathways were globally always downregulated under the higher light level, regardless of the light treatment comparison (0.1 lux *vs* controls, 5 lux *vs* controls or 5 lux *vs* 0.1 lux) at night or daytime (S2 Table). In contrast, among the differentially expressed genes mentioned previously (FDR < 0.05 and |LFC| > 2), we did not find any gene related to circadian rhythm, *i.e.,* clock genes or genes coding for enzymes involved in melatonin synthesis, which we presumed to be affected by ALAN. Therefore, no GO terms involved in circadian rhythms were found to be enriched (S3 Table).

## Discussion

Our study of 30 transcriptomes from *B. bufo* tadpoles exposed to light treatments during two thirds of larval development demonstrated that ALAN induced a strong disruption of gene expression both at night and daytime. In addition, we clearly showed that most of the differentially expressed genes when tadpoles experiment ALAN were downregulated, leading to changes in many physiological pathways, including their innate immune system and their lipid metabolism.

At night, ALAN induced strong differential gene expression compared to controls. More unexpectedly, we showed noticeable differential gene expression during the daytime between tadpoles exposed to ALAN and controls, but only at our high ALAN level (5 lux). To our knowledge, all published studies to date at the molecular level have always focused on variations in gene expression occurring at night during ALAN exposure but never during the day, when light levels are similar between exposed individuals and controls (Honnen et al., 2016; Rosenberg et al., 2019). Our results suggest that these alterations in gene expression are a direct long-lasting result of exposure to ALAN during the night-time, and an indirect effect of physiological disorders impacting gene expression during the daytime. Moreover, our results showed that ALAN has a rather monotonic dose-dependent effect on molecular parameters. Indeed, the higher the ALAN illuminance, the greater the deregulation of transcriptome expression. In addition, while our low ALAN intensity (0.1 lux) affects gene expression only during night-time, at our high ALAN intensity (5 lux) gene expression is altered both at night-time and daytime. Thus, it is necessary to consider not only the effects at night when ALAN is present but also the residual disturbances still occurring during the day in the absence of this stressor. In particular, molecular impacts were obtained with very low illuminance corresponding to ecological light pollution measured *in natura*. The 0.1 lux intensity is relevant to the skyglow level recorded in rural areas (Gaston et al., 2013). We observed significant molecular effects of ALAN in one species, the common toad, which is known to tolerate proximity to humans well, as the common toad is often found near human settlement, suggesting that this observation is not very reassuring for other species in this group, which is globally in a poor conservation status worldwide (The IUCN red list of threatened species).

Analysis of differentially expressed genes and GO term enrichment showed a reduced expression of genes involved in innate immune system functioning during both night-time and daytime. GO terms linked to antibacterial humoral response, inflammatory response, defence response, C5a anaphylatoxin chemotactic receptor binding, regulation of membrane attack complex, complement-dependent cytotoxicity and complement activation were significantly enriched in differentially expressed genes. Genes downregulated at night and daytime under ALAN were, for instance, coding for complement C3, complement C3 alpha chain and complement factor H. In vertebrates, the activation of the complement system, a key component of innate immunity, acts by promoting the inflammatory response to eliminate microorganisms (Demas et al., 2011). The complement is also involved in the lysis of foreign cells through the formation of membrane attack complexes and the mediation of phagocytosis (Demas et al., 2011; Juul-Madsen et al., 2014). A nocturnal downregulation of genes coding for peptidoglycan-recognition proteins was also observed under ALAN. These proteins play a key role in innate immunity, as they belong to a group of highly conserved pattern recognition receptors involved in the recognition of the bacterial peptidoglycan wall (Dziarski & Gupta, 2006). It would, therefore, be relevant to investigate whether this downregulation of genes involved in immune function results in depressed immunocompetence in tadpoles exposed to ALAN. Several studies have indeed shown a modification in the immune capacity of organisms in the presence of ALAN, even if these effects were complex and sometimes contradictory. For instance, ALAN reduced the bactericidal activity of blood plasma and cell-mediated immune function in the Siberian hamster, *Phodopus sungorus* (Bedrosian et al., 2011), while those parameters were increased by ALAN in the king quail, *Excalfactoria chinensis* (Saini et al., 2019). This suggests that the immune response following ALAN exposure is not uniform across species, and such heterogeneity is possibly related to light intensity, duration of treatment and potential developmental window of sensitivity. ALAN effects on the immune system have indeed been shown to be stage-dependent in birds (Ouyang et al., 2017; Raap, Casasole, Pinxten, et al., 2016). Adult great tits, *Parus major*, exposed to 8.2 lux show no change in haptoglobin concentration (Ouyang et al., 2017), characteristic of the nonspecific immune response, whereas in chicks of the same species, exposure to 5 lux induces an increase in this parameter (Raap, Casasole, Pinxten, et al., 2016), suggesting increased early sensitivity. In vertebrates, the innate immune system provides rapid and nonspecific protection against a variety of immune challenges (Demas et al., 2011). In the wild, amphibians encounter a variety of immune challenges, particularly due to the presence parasites, pathogens and of two infectious fungal species, *Batrachochytrium dendrobatidis* and *Batrachochytrium salamandrivorans* (Grogan et al., 2018), representing a major cause of the decline in amphibians worldwide. Although infection with infectious fungal species is generally not lethal before metamorphosis, infected tadpoles may exhibit sublethal effects on growth and development (Blaustein et al., 2005). Thus, by reducing the innate immune response, exposure to light pollution *in natura* would probably impair the capacity of tadpoles to overcome immune challenges, thus altering their survival. A similar experimental study conducted on the impact of traffic noise on European treefrogs, *Hyla arborea*, emphasized a strong decrease in the immune response in adults exposed to noise (Troïanowski et al., 2017). In contrast, in adults of four amphibian species, urban habitats that include light pollution were not associated with impaired cutaneous immune activity, as estimated by delayed-type hypersensitivity after phytohaemagglutinin injection *in vivo* (Iglesias-Carrasco et al., 2017). The present data may therefore suggest a particular sensitivity of the immune system to ALAN at the tadpole stage.

GO terms linked to lipid metabolism and transport were another important physiological pathway affected by ALAN. For the first time, we demonstrated that this function was affected at realistic level of ALAN (0.1 lux). In an experimental study conducted with an ALAN level of 2 lux, an impaired balance of lipid biosynthetic pathways has been demonstrated in rodents (Okuliarova et al., 2020) and in mosquitoes but exposed to 300 lux at night (Honnen et al., 2016). ALAN-related disruption of circadian rhythmicity through a modification of the expression of clock genes has been suggested to be involved in this impairment (Fonken & Nelson, 2014; Rumanova et al., 2020). As in the majority of organisms, in amphibians, lipid metabolic activity occurs mainly in the liver and its regulation is based on the balance between the synthesis of fatty acids and fat catabolism (Sheridan, 1994). A modification of the expression of genes related to lipid metabolism and transport in the presence of ALAN, as observed at 0.1 and 5 lux both during night-time and daytime, is likely to affect body lipid homeostasis. Such disruption of lipid metabolism would likely limit the availability and use of lipid resources, and thus be of particular concern in developing individuals. Indeed, lipids have the potential of serving as a source of metabolic energy (Sawant & Varute, 1973). They also seems to form an important constituent of the tadpole bodies, as total lipids are known to increase during the phase of tissue and organ growth involving cell divisions and differentiation as happens in the larval growth stages and prometamorphosis (Sawant & Varute, 1973). During metamorphosis, when histolytic events appear, *e.g.,* internal gills degeneration, tail regression, and when tadpoles do not feed, *i.e.,* no exogenous source of energy is available and full reliance on endogenous source, anuran tadpoles seem to make use of lipids that have accumulated during the growth period (Sawant & Varute, 1973). Thus, studies of the consequences of ALAN on the physical activity and growth of juveniles in relation to metabolism should further investigate this issue.

The majority of ALAN studies have suggested that light at night affects circadian rhythms through modifications of clock gene regulation and melatonin synthesis. However, when comparing our transcriptomes at night or daytime, neither clock genes (*e.g.,* Clock, Period, Bmal 1, Cryptochrome) nor melatonin-related genes (*e.g.,* genes encoding enzymes catalysing melatonin synthesis) were differentially expressed between light treatments. Present results contrast with those obtained by others at 5 lux, where melatonin plasma levels were reduced by more than a quarter in European sea bass, *Dicentrarchus labrax* (Bayarri et al., 2002), in great tit, *P. major* (de Jong et al., 2016) or in zebra finch, *Taeniopygia guttata* (Batra et al., 2019). An absence of changes in melatonin-related genes could possibly be related to the fact that enzyme expression did not reflect melatonin synthesis in *B. Bufo* tadpoles. Indeed, a positive correlation between retinal aralkylamine-N-acetyltransferase (AANAT) activity, an enzyme catalysing melatonin synthesis, and retinal melatonin levels during the light/dark cycle was not observed in adult European green toads, *Bufo viridis* (Serino et al., 1993). A positive correlation was however reported in adult green frogs, *Pelophylax esculenta* (d’Istria et al., 1994; Serino et al., 1993). Alternatively, because of the low levels of ALAN intensities used in our study, this could have resulted in discrete changes in the transcription of these genes that were not detected by our large-scale analysis. Clarifying this point warrants further investigations with more specific approaches.

In conclusion, the present study revealed that ecologically relevant illuminances at night mimicking anthropic emerging pollution markedly affected the global gene expression pattern in tadpoles of common toads. Transcriptomic changes suggested that ALAN would result in depressed capacity to overcome immune environmental challenges, thus potentially impairing tadpole survival. The impacts of ALAN are multifarious, and we only slowly disentangle its multiple negative effects on organisms, populations and ecosystems. Present data advocate the need for integrative studies with molecular approaches to identify and characterize pathways that may link physiological and behavioural disruption caused by light at night and potential health and *fitness* consequences.

## Material and Methods

### Animal collection and housing conditions

Nine fragments of approximately 20 eggs each were randomly sampled from a clutch of common toads in March at the beginning of the breeding season in La Burbanche, France (45°N, 5°E). This site was chosen for its low levels of ALAN regardless of weather conditions and lunar phase (≤ 0.01 lux). Upon arrival at the animal care facility (EcoAquatron, University of Lyon), fragments were placed individually in an aquarium (32 cm × 17.5 cm × 18.5 cm) containing 7.4 L of dechlorinated and bubbled water. From stage 25 of Gosner, the stage at which independent feeding is possible (Gosner, 1960), to the end of the experiment, tadpoles were fed every two days with boiled organic green salad. Each week, one-third of the water of the aquarium was renewed with dechlorinated and oxygenated water. Ambient temperature and relative humidity were kept constant during the whole experiment at 16.1 ± 0.9°C and 60.0 ± 11.2%, respectively.

### Light treatments

Aquariums were immediately randomly assigned to one of the three light treatments (3 aquariums per treatment) and exposed to their respective light treatment at night. During the daytime, illuminance provided by light tubes (Philips Master TL-D 58 W/865 and Exo Terra Repti Glo 2.0, 40 W T8) for all aquariums was 1995 ± 257 lux (mean ± SEM). During night-time, we used white light-emitting diode (LED) ribbons (white cold Light Plus, 6000-6500 K, 14 W, 60 LED/m) (Touzot et al., 2020) to reproduce artificial light by night. White LEDs were chosen because they are increasingly used for street lighting worldwide (Falchi et al., 2016). An LED ribbon of 95 cm (57 LED) combined with a light diffuser was suspended horizontally 41.5 cm above the bottom of the aquariums. Each LED ribbon was connected to a dimmer (manual dimmer, 12 V max, 8 A) and a laboratory power supply (15 V/DC max, 3 A), which allowed illuminance to be set for each light treatment. The aquariums of one light treatment were isolated from the others with tarpaulins to avoid light spilling over from another treatment. Tadpoles were exposed to the same photoperiod as the natural photoperiod at the date of the experiment. At night, the control group was exposed to 0.01 lux, corresponding to the illuminance of a sky under clear conditions with a quarter moon (Gaston et al., 2013). The first experimental group was exposed to 0.1 lux, corresponding to the illuminance of urban skyglow (Gaston et al., 2013). The second experimental group was exposed to 5 lux, which corresponds to the light level of a residential street (Gaston et al., 2013). The daylight and 5 lux illuminances were measured with a luxmeter (Illuminance meter T-10A, Konica Minolta). Illuminances for the 0.1 lux and control illuminances were set using a highly sensitive light meter (Sky Quality Meter SQM-L, Unihedron). Consequently, to compare illuminances, SQM-measured values were converted into lux according to the relationship curve between SQM and lux measurements (Touzot et al., 2020), as lux is the main unit used in ALAN studies (Longcore & Rich, 2004). Illuminances were measured at the water surface of each aquarium (24 cm from the LED) and checked every week (0.007 ± 0.001 lux for the control group, 0.02 ± 0.004 lux for the 0.1-lux group and 4.09 ± 1.30 lux for the 5-lux-group, mean ± SEM).

### Experiment

When tadpoles reached stage 31 of Gosner (Gosner, 1960), corresponding to 27 days of exposure to ALAN, 5 tadpoles of each treatment were randomly sampled during daytime (13:00) or night-time (01:00) (random sampling on the 3 aquariums of each treatment). Six hours prior to sampling, tadpoles were fasted so that the nutritional status was the same for all animals and did not bias gene expression. Development stage 31 was chosen, as it is an easily identifiable stage, and the foot is paddle-shaped at this stage only (Gosner, 1960). Development stage 31 also allowed long-term exposure to ALAN, while being distant enough from metamorphosis, *i.e.,* stage 46 of Gosner, the ultimate stages of development during which thyroid hormones are strongly expressed and involve many functional and morphological changes in the organisms. Tadpoles were individually sampled and immediately placed in liquid nitrogen and then stored at −80°C until molecular analyses. Although all individuals were at the same Gosner stage, the tail size differed between individuals, as the tadpole tail was growing. Thus, the tadpole tail was removed before molecular analyses to limit the effect of tail growth on gene expression. Although gene expression can differ among tissues, we chose to work on the body without distinguishing tissues to investigate the global effects of ALAN across as broad a range of molecular processes as possible.

### Molecular analyses

Total RNA was extracted by adding TRI Reagent (Molecular Research Center MRC, TR118) to the 30 samples and by homogenizing tissues on ice with a tissue lyser (Retsch, MM200). All remaining steps were carried out according to the manufacturers’ protocols (Molecular Research Center MRC, TR118). Then, the RNA was treated with Turbo DNase enzyme (Turbo DNA free kit, Invitrogen, AM1907) and assayed by fluorescence using a Qubit fluorometer (Qubit® RNA HS Assay Kits Molecular Probes, Invitrogen, Q32855). Library construction was carried out using the NEBNext Ultra II RNA Library Prep Kit for Illumina (Biolabs New England, NEB E7770, E7490, E7600). Libraries were randomly sequenced on three lanes with an Illumina® HiSeq4000™ machine on the GenomEast platform hosted at the IGBMC (GenomEast Platform – Institut de Génétique et de la Biologie Moléculaire et Cellulaire IGBMC, Illkirch, France), resulting in single-end samples with reads 50 bp long.

### *B. bufo de novo* transcriptome assembly

We built a transcriptome assembly from the sequenced RNA samples (S1 Appendix) using Trinity software (version 2.5.1) (Haas et al., 2013) with default parameters. All samples were processed in a single run with the Trinity default read normalization procedure to improve the sensitivity of transcript detection. The resulting assembly was assessed using Busco software (version 3.0.2) (Seppey et al., 2019). We ran Busco using the Tetrapoda dataset (available at https://busco-archive.ezlab.org/v2/datasets/tetrapoda_odb9.tar.gz) and with an E-value threshold of 0.01 (--evalue option). Protein sequences and valid coding sequences (CDSs) were derived from assembled transcripts using the Transdecoder program (version 3.0.1) (Haas et al., 2013). We thus obtained a filtered set of transcripts that was subsequently fed to the Diamond program (Buchfink et al., 2015) with default arguments to discover homologies with genes from the reference genome of *Xenopus tropicalis*, and proteins from the UniProt/Swiss-Prot database. In addition, the CDSs were annotated using PFAM (El-Gebali et al., 2019) domains using the hmmsearch program (Wheeler & Eddy, 2013) with default arguments.

### Differential gene expression analysis

Expression levels were quantified using Kallisto (Bray et al., 2016) with options −1200 −s 20 for single-end samples. We subsequently used rounded effective counts as input for differential gene expression, which was performed using the *DESeq2* package (Love et al., 2014). We specified light treatment, timepoint and their interaction in the model (design formula: “design = ~ timepoint + light treatment + timepoint * light treatment). The *DESeq2* package provides differential gene expression by use of negative binomial distribution and shrinkage estimation for dispersions and fold changes to improve the stability and interpretability of estimates (Love et al., 2014). P-Values were used to test the significance of the false rate discovery (FDR) using the Benjamini and Hochberg procedure. Genes with an FDR < 0.05 were considered significantly differentially expressed. We also reported numbers of genes with an absolute Log2 Fold Change (|LFC|) greater than 1 and 2. Moreover, we ran a PCA using the *ggplot2* package for the 30 samples. Proportional Venn diagrams were constructed using the *eulerr* package and bar plots were constructed using the *ggplot2* package.

### Gene enrichment analysis

Functional annotations and overrepresentation of GO terms were analysed using the *topGO* package (Alexa & Rahnenfuhrer, 2020). Gene enrichment analyses were performed using a hypergeometric test for overrepresentation of GO terms (Fisher test) from the differentially expressed gene set (FDR < 0.05 and |LFC| > 1). Again, P-values were used to test FDR significance using the Benjamini and Hochberg procedure, and GO terms with FDR < 0.01 and odds-ratio > 1 were regarded as significantly different. Then, GO term redundancy was removed first using REVIGO (Supek et al., 2011) and second by manual simplification as GO terms were still redundant. The second simplification occurred when two GO terms were parents/descendants and were significantly enriched with the same number of annotated and significantly differentially expressed genes. We evaluated GO terms from the biological process, molecular function and cellular component domains.

The analyses of differential gene expression and gene enrichment, and graphics were performed with R software (v3.6.0) (*R Core Team*, 2018).

### Ethical note

The sampling of common toad clutch fragments was authorized by the Préfecture de l’Ain (DDPP01-18-271) and by the French government (*APAFIS*#27760-2020102115402767) in accordance with the ethical committee of Lyon 1 University. The animal care structure “EcoAquatron” (University of Lyon) received an agreement of veterinary services (approval DSV 692661201). At the end of the experiment, all the remaining tadpoles were released on the capture site.

### Data availability

Transcriptomes reads have been deposited on the Gene Expression Omnibus (GEO) of the Sequence Read Archive (SRA) and are available under accession numbers from XXXXXXXX to XXXXXXXX in the study XXXXXXXXXX. *B. bufo de novo* transcriptome has been deposited on the Sequence Read Archive (SRA) and are available under accession number XXXXXXXX.

## Acknowledgements

We thank A. Clair, J. Ulmann and L. Averty for their technical assistance in the EcoAquatron and M. Sémon for her advice on the experimental design and data analysis.

## Author contributions

M. Touzot: Conceptualization, Formal Analysis, Investigation, Writing – Original Draft Preparation.

T. Lefebure: Data Curation, Formal Analysis, Writing – Review and Editing.

T. Lengagne and J. Secondi: Funding Acquisition, Writing – Review and Editing.

A. Dumet and L. Konecny-Dupre: Investigation.

P. Veber and V. Navratil: Data Curation, Writing – Review and Editing.

C. Duchamp: Writing – Review and Editing.

N. Mondy: Conceptualization, Investigation, Writing – Review and Editing.

## Supporting information

**S1 Table:** Enrichment level of the 54 most enriched GO terms

**S2 Table:** The top 10 most overrepresented GO terms

**S3 Table:** Enrichment level of GO terms related to circadian rhythm

**S1 Appendix:** Sequencing and assembly of *B. bufo de novo* transcriptome

## References

Alexa, A., & Rahnenfuhrer, J. (2020). TopGO: Enrichment Analysis for Gene Ontology. https://doi.org/10.18129/B9.BIOC.TOPGO

Batra, T., Malik, I., & Kumar, V. (2019). Illuminated night alters behaviour and negatively affects physiology and metabolism in diurnal zebra finches. Environmental Pollution, 254, 112916. https://doi.org/10.1016/j.envpol.2019.07.084

Bayarri, M. J., Madrid JA, & SaÂnchez-VaÂzquez FJ. (2002). Influence of light intensity, spectrum and orientation on sea bass plasma and ocular melatonin. Journal of Pineal Research, 32, 34–40. https://doi.org/10.1034/j.1600-079x.2002.10806.x

Bedrosian, T. A., Fonken, L. K., Walton, J. C., & Nelson, R. J. (2011). Chronic exposure to dim light at night suppresses immune responses in Siberian hamsters. Biology Letters, 7(3), 468–471. https://doi.org/10.1098/rsbl.2010.1108

Bedrosian, T. A., Galan, A., Vaughn, C. A., Weil, Z. M., & Nelson, R. J. (2013). Light at Night Alters Daily Patterns of Cortisol and Clock Proteins in Female Siberian Hamsters. Journal of Neuroendocrinology, 25(6), 590–596. https://doi.org/10.1111/jne.12036

Beebee, T. J. C. (1979). Habitats of the British amphibians (2) : Suburban parks and gardens. Biological Conservation, 15(4), 241–257. https://doi.org/10.1016/0006-3207(79)90046-6

Blaustein, A. R., Romansic, J. M., Scheessele, E. A., Han, B. A., Pessier, A. P., & Longcore, J. E. (2005). Interspecific Variation in Susceptibility of Frog Tadpoles to the Pathogenic Fungus Batrachochytrium dendrobatidis. Conservation Biology, 19(5), 1460–1468. https://doi.org/10.1111/j.1523-1739.2005.00195.x

Bray, N. L., Pimentel, H., Melsted, P., & Pachter, L. (2016). Near-optimal probabilistic RNA-seq quantification. Nature Biotechnology, 34(5), 525–527. https://doi.org/10.1038/nbt.3519

Brüning, A., Kloas, W., Preuer, T., & Hölker, F. (2018). Influence of artificially induced light pollution on the hormone system of two common fish species, perch and roach, in a rural habitat. Conservation Physiology, 6(1). https://doi.org/10.1093/conphys/coy016

Buchfink, B., Xie, C., & Huson, D. H. (2015). Fast and sensitive protein alignment using DIAMOND. Nature Methods, 12(1), 59–60. https://doi.org/10.1038/nmeth.3176

de Jong, M., Jeninga, L., Ouyang, J. Q., van Oers, K., Spoelstra, K., & Visser, M. E. (2016). Dose-dependent responses of avian daily rhythms to artificial light at night. Physiology & Behavior, 155, 172–179. https://doi.org/10.1016/j.physbeh.2015.12.012

Demas, G. E., Zysling, D. A., Beechler, B. R., Muehlenbein, M. P., & French, S. S. (2011). Beyond phytohaemagglutinin : Assessing vertebrate immune function across ecological contexts: Assessing vertebrate immune function across ecological contexts. Journal of Animal Ecology, 80(4), 710–730. https://doi.org/10.1111/j.1365-2656.2011.01813.x

d’Istria, M., Monteleone, P., Serino, I., & Chieffi, G. (1994). Seasonal variations in the daily rhythm of melatonin and NAT activity in the Harderian gland, retina, pineal gland, and serum of the green frog, Rana esculenta. General and comparative endocrinology, 96(1), 6–11.

Dominoni, D. M., de Jong, M., van Oers, K., O’Shaughnessy, P., Blackburn, G., Atema, E., Mateman, C. A., D’Amelio, P. B., Trost, L., Bellingham, M., Clark, J., Visser, M. E., & Helm, B. (2020). Artificial light at night shifts the circadian system but still leads to physiological disruption in a wild bird [Preprint]. Zoology. https://doi.org/10.1101/2020.12.18.423473

Dudgeon, D., Arthington, A. H., Gessner, M. O., Kawabata, Z.-I., Knowler, D. J., Lévêque, C., Naiman, R. J., Prieur-Richard, A.-H., Soto, D., Stiassny, M. L. J., & Sullivan, C. A. (2006). Freshwater biodiversity : Importance, threats, status and conservation challenges. Biological Reviews, 81(02), 163–182. https://doi.org/10.1017/S1464793105006950

Dziarski, R., & Gupta, D. (2006). The peptidoglycan recognition proteins (PGRPs). Genome Biology, 7(8), 232. https://doi.org/10.1186/gb-2006-7-8-232

El-Gebali, S., Mistry, J., Bateman, A., Eddy, S. R., Luciani, A., Potter, S. C., Qureshi, M., Richardson, L. J., Salazar, G. A., Smart, A., Sonnhammer, E. L. L., Hirsh, L., Paladin, L., Piovesan, D., Tosatto, S. C. E., & Finn, R. D. (2019). The Pfam protein families database in 2019. Nucleic Acids Research, 47(D1), D427‑D432. https://doi.org/10.1093/nar/gky995

Falchi, F., Cinzano, P., Duriscoe, D., Kyba, C. C. M., Elvidge, C. D., Baugh, K., Portnov, B. A., Rybnikova, N. A., & Furgoni, R. (2016). The new world atlas of artificial night sky brightness. Science Advances, 2(6), e1600377. https://doi.org/10.1126/sciadv.1600377

Fonken, L. K., Aubrecht, T. G., Meléndez-Fernández, O. H., Weil, Z. M., & Nelson, R. J. (2013). Dim light at night disrupts molecular circadian rhythms and increases body weight. Journal of Biological Rhythms, 28(4), 262–271. https://doi.org/10.1177/0748730413493862

Fonken, L. K., & Nelson, R. J. (2014). The Effects of Light at Night on Circadian Clocks and Metabolism. Endocrine Reviews, 35(4), 648–670. https://doi.org/10.1210/er.2013-1051

Gaston, K. J., Bennie, J., Davies, T. W., & Hopkins, J. (2013). The ecological impacts of nighttime light pollution : A mechanistic appraisal: Nighttime light pollution. Biological Reviews, 88(4), 912–927. https://doi.org/10.1111/brv.12036

Gaston, K. J., Visser, M. E., & Hölker, F. (2015). The biological impacts of artificial light at night : The research challenge. Philosophical Transactions of the Royal Society B: Biological Sciences, 370(1667), 20140133. https://doi.org/10.1098/rstb.2014.0133

Gosner, K. L. (1960). A Simplified Table for Staging Anuran Embryos and Larvae with Notes on Identification. Herpetologica, 16(3), 183–190.

Grogan, L. F., Robert, J., Berger, L., Skerratt, L. F., Scheele, B. C., Castley, J. G., Newell, D. A., & McCallum, H. I. (2018). Review of the Amphibian Immune Response to Chytridiomycosis, and Future Directions. Frontiers in Immunology, 9, 2536. https://doi.org/10.3389/fimmu.2018.02536

Grubisic, M. (2018). Waters under Artificial Lights : Does Light Pollution Matter for Aquatic Primary Producers? Limnology and Oceanography Bulletin, 27(3), 76–81. https://doi.org/10.1002/lob.10254

Grubisic, M., Haim, A., Bhusal, P., Dominoni, D. M., Gabriel, K. M. A., Jechow, A., Kupprat, F., Lerner, A., Marchant, P., Riley, W., Stebelova, K., van Grunsven, R. H. A., Zeman, M., Zubidat, A. E., & Hölker, F. (2019). Light Pollution, Circadian Photoreception, and Melatonin in Vertebrates. Sustainability, 11(22), 6400. https://doi.org/10.3390/su11226400

Guetté, A., Godet, L., Juigner, M., & Robin, M. (2018). Worldwide increase in Artificial Light At Night around protected areas and within biodiversity hotspots. Biological Conservation, 223, 97–103. https://doi.org/10.1016/j.biocon.2018.04.018

Haas, B. J., Papanicolaou, A., Yassour, M., Grabherr, M., Blood, P. D., Bowden, J., Couger, M. B., Eccles, D., Li, B., Lieber, M., MacManes, M. D., Ott, M., Orvis, J., Pochet, N., Strozzi, F., Weeks, N., Westerman, R., William, T., Dewey, C. N., … Regev, A. (2013). De novo transcript sequence reconstruction from RNA-seq using the Trinity platform for reference generation and analysis. Nature Protocols, 8(8), 1494–1512. https://doi.org/10.1038/nprot.2013.084

Haim, A., & Portnov, B. A. (2013). Light Pollution as a New Risk Factor for Human Breast and Prostate Cancers. Springer Netherlands. https://doi.org/10.1007/978-94-007-6220-6

Hölker, F., Wolter, C., Perkin, E. K., & Tockner, K. (2010). Light pollution as a biodiversity threat. Trends in Ecology & Evolution, 25(12), 681–682. https://doi.org/10.1016/j.tree.2010.09.007

Honnen, A.-C., Johnston, P. R., & Monaghan, M. T. (2016). Sex-specific gene expression in the mosquito Culex pipiens f. Molestus in response to artificial light at night. BMC Genomics, 17(1), 22. https://doi.org/10.1186/s12864-015-2336-0

Honnen, A.-C., Kypke, J. L., Hölker, F., & Monaghan, M. T. (2019). Artificial Light at Night Influences Clock-Gene Expression, Activity, and Fecundity in the Mosquito Culex pipiens f. Molestus. Sustainability, 11(22), 6220. https://doi.org/10.3390/su11226220

Iglesias-Carrasco, M., Martín, J., & Cabido, C. (2017). Urban habitats can affect body size and body condition but not immune response in amphibians. Urban Ecosystems, 20(6), 1331–1338. https://doi.org/10.1007/s11252-017-0685-y25

Juul-Madsen, H. R., Viertlböeck, B., Härtle, S., Smith, A. L., & Göbel, T. W. (2014). Innate Immune Responses. In Avian Immunology (p. 121‑147). Elsevier. https://doi.org/10.1016/B978-0-12-396965-1.00007-8

Kupprat, F., Hölker, F., Knopf, K., Preuer, T., & Kloas, W. (2021). Innate immunity, oxidative stress and body indices of Eurasian perch *Perca fluviatilis* after two weeks of exposure to artificial light at night. Journal of Fish Biology, jfb.14703. https://doi.org/10.1111/jfb.14703

Latchem, E., Madliger, C. L., Abrams, A. E. I., & Cooke, S. J. (2021). Does Artificial Light at Night Alter the Subsequent Diurnal Behavior of a Teleost Fish? Water, Air, & Soil Pollution, 232(2), 71. https://doi.org/10.1007/s11270-021-05023-4

Longcore, T., & Rich, C. (2004). Ecological light pollution. Frontiers in Ecology and the Environment, 2(4), 191–198. https://doi.org/10.1890/1540-9295(2004)002[0191:elp]2.0.co;2

Love, M. I., Huber, W., & Anders, S. (2014). Moderated estimation of fold change and dispersion for RNA-seq data with DESeq2. Genome Biology, 15(12), 550. https://doi.org/10.1101/002832

Luginbuhl, C. B., Boley, P. A., & Davis, D. R. (2014). The impact of light source spectral power distribution on sky glow. Journal of Quantitative Spectroscopy and Radiative Transfer, 139, 21–26. https://doi.org/10.1016/j.jqsrt.2013.12.004

Okuliarova, M., Rumanova, V. S., Stebelova, K., & Zeman, M. (2020). Dim Light at Night Disturbs Molecular Pathways of Lipid Metabolism. International Journal of Molecular Sciences, 21(18), 6919. https://doi.org/10.3390/ijms21186919

Ouyang, J. Q., de Jong, M., van Grunsven, R. H. A., Matson, K. D., Haussmann, M. F., Meerlo, P., Visser, M. E., & Spoelstra, K. (2017). Restless roosts : Light pollution affects behavior, sleep, and physiology in a free-living songbird. Global Change Biology, 23(11), 4987–4994. https://doi.org/10.1111/gcb.13756

R Core Team. (2018). https://www.R-project.org/

Raap, T., Casasole, G., Costantini, D., AbdElgawad, H., Asard, H., Pinxten, R., & Eens, M. (2016). Artificial light at night affects body mass but not oxidative status in free-living nestling songbirds : An experimental study. Scientific Reports, 6, 35626. https://doi.org/10.1038/srep35626

Raap, T., Casasole, G., Pinxten, R., & Eens, M. (2016). Early life exposure to artificial light at night affects the physiological condition : An experimental study on the ecophysiology of free-living nestling songbirds. Environmental Pollution, 218, 909–914. https://doi.org/10.1016/j.envpol.2016.08.024

Reid, A. J., Carlson, A. K., Creed, I. F., Eliason, E. J., Gell, P. A., Johnson, P. T. J., Kidd, K. A., MacCormack, T. J., Olden, J. D., Ormerod, S. J., Smol, J. P., Taylor, W. W., Tockner, K., Vermaire, J. C., Dudgeon, D., & Cooke, S. J. (2019). Emerging threats and persistent conservation challenges for freshwater biodiversity. Biological Reviews, 94(3), 849–873. https://doi.org/10.1111/brv.12480

Reiter, R. J., Tan, D.-X., & Fuentes-Broto, L. (2010). Melatonin : A Multitasking Molecule. Progress in Brain Research, 181, 127–151. https://doi.org/10.1016/S0079-6123(08)81008-4

Robert, K. A., Lesku, J. A., Partecke, J., & Chambers, B. (2015). Artificial light at night desynchronizes strictly seasonal reproduction in a wild mammal. Proc Biol Sci, 282(1816), 20151745. https://doi.org/10.1098/rspb.2015.1745

Rosenberg, Y., Doniger, T., & Levy, O. (2019). Sustainability of coral reefs are affected by ecological light pollution in the Gulf of Aqaba/Eilat. Communications Biology, 2(1), 289. https://doi.org/10.1038/s42003-019-0548-6

Rumanova, V. S., Okuliarova, M., & Zeman, M. (2020). Differential Effects of Constant Light and Dim Light at Night on the Circadian Control of Metabolism and Behavior. International Journal of Molecular Sciences, 21(15), 5478. https://doi.org/10.3390/ijms21155478

Saini, C., Hutton, P., Gao, S., Simpson, R. K., Giraudeau, M., Sepp, T., Webb, E., & McGraw, K. J. (2019). Exposure to artificial light at night increases innate immune activity during development in a precocial bird. Comparative Biochemistry and Physiology Part A: Molecular & Integrative Physiology, 233, 84–88. https://doi.org/10.1016/j.cbpa.2019.04.002

Sawant, V. A., & Varute, A. T. (1973). Lipid changes in the tadpoles of Rana tigrina during growth and metamorphosis. Comparative Biochemistry and Physiology Part B: Comparative Biochemistry, 44(3), 729–750. https://doi.org/10.1016/0305-0491(73)90223-X

Secondi, J., Davranche, A., Théry, M., Mondy, N., & Lengagne, T. (2019). Assessing the effects of artificial light at night on biodiversity across latitude – Current knowledge gaps. Global Ecology and Biogeography, 29(3), 404–419. https://doi.org/10.1111/geb.13037

Secondi, J., Dupont, V., Davranche, A., Mondy, N., Lengagne, T., & Théry, M. (2017). Variability of surface and underwater nocturnal spectral irradiance with the presence of clouds in urban and peri-urban wetlands. PLOS ONE, 12(11), e0186808. https://doi.org/10.1371/journal.pone.0186808

Seppey, M., Manni, M., & Zdobnov, E. M. (2019). BUSCO : Assessing Genome Assembly and Annotation Completeness. In M. Kollmar (Éd.), Gene Prediction (Vol. 1962, p. 227‑245). Springer New York. https://doi.org/10.1007/978-1-4939-9173-0_14

Serino, I., D’Istria, M., & Monteleone, P. (1993). A comparative study of melatonin production in the retina, pineal gland and Harderian gland of Bufo viridis and Rana esculenta. Comparative Biochemistry and Physiology Part C: Pharmacology, Toxicology and Endocrinology, 106(1), 189–193. https://doi.org/10.1016/0742-8413(93)90271-L

Sheridan, M. A. (1994). Regulation of lipid metabolism in poikilothermic vertebrates. Comparative Biochemistry and Physiology Part B: Comparative Biochemistry, 107(4), 495–508. https://doi.org/10.1016/0305-0491(94)90176-7

Supek, F., Bošnjak, M., Škunca, N., & Šmuc, T. (2011). REVIGO Summarizes and Visualizes Long Lists of Gene Ontology Terms. PLoS ONE, 6(7), e21800. https://doi.org/10.1371/journal.pone.0021800

The IUCN red list of threatened species. (s. d.). International Union for Conservation of Nature, IUCN, https://www.iucnredlist.org/. Consulté 18 mai 2020, à l’adresse https://www.iucnredlist.org/.

Touzot, M., Lengagne, T., Secondi, J., Desouhant, E., Théry, M., Dumet, A., Duchamp, C., & Mondy, N. (2020). Artificial light at night alters the sexual behaviour and fertilisation success of the common toad. Environmental Pollution, 259, 113883. https://doi.org/10.1016/j.envpol.2019.113883

Troïanowski, M., Mondy, N., Dumet, A., Arcanjo, C., & Lengagne, T. (2017). Effects of traffic noise on tree frog stress levels, immunity, and color signaling : Noise Consequences on Tree Frogs. Conservation Biology, 31(5), 1132–1140. https://doi.org/10.1111/cobi.12893

Wheeler, T. J., & Eddy, S. R. (2013). nhmmer : DNA homology search with profile HMMs. Bioinformatics, 29(19), 2487–2489. https://doi.org/10.1093/bioinformatics/btt403

Wyse, C. A., Selman, C., Page, M. M., Coogan, A. N., & Hazlerigg, D. G. (2011). Circadian desynchrony and metabolic dysfunction; did light pollution make us fat? Medical Hypotheses, 77(6), 1139–1144. https://doi.org/10.1016/j.mehy.2011.09.023

Zhao, D., Yu, Y., Shen, Y., Liu, Q., Zhao, Z., Sharma, R., & Reiter, R. J. (2019). Melatonin Synthesis and Function : Evolutionary History in Animals and Plants. Frontiers in Endocrinology, 10, 249. https://doi.org/10.3389/fendo.2019.0024

