## supporting_information for "Large-scale deregulation of gene expression by artificial light at night in tadpoles of common toads"

**S1 Table:** Enrichment level of the 54 most enriched GO terms (FDR < 0.01 and odds-ratio > 10 for at least one light treatment comparison) for each light treatment comparisons at either night-time or daytime.

| | | At night-time | | | At daytime | | |
|---|---|---|---|---|---|---|---|
| | | **0.1 lux *vs* control** | **5 lux *vs* control** | **5 lux *vs* 0.1 lux** | **0.1 lux *vs* control** | **5 lux *vs* control** | **5 lux *vs* 0.1 lux** |
| **GO** | **Annotation** | **Odds-ratio (FDR)** | **Odds-ratio (FDR)** | **Odds-ratio (FDR)** | **Odds-ratio (FDR)** | **Odds-ratio (FDR)** | **Odds-ratio (FDR)** |
| GO:0001794 | Type IIa hypersensitivity | 0.00 (1.10$^{1}$) | 28.57 (2.10$^{-3}$) | 150.00 (3.10$^{-4}$) | 0.00 (1.10$^{1}$) | 26.67 (4.10$^{-4}$) | 16.67 (2.10$^{-2}$) |
| GO:0001868 | Regulation of complement activation lectin pathway | 0.00 (1.10$^{1}$) | 4.88 (8.10$^{-1}$) | 28.57 (1.10$^{-1}$) | 0.00 (1.10$^{1}$) | 11.36 (4.10$^{-3}$) | 9.43 (7.10$^{-3}$) |
| GO:0001905 | Activation of membrane attack complex | 0.00 (1.10$^{1}$) | 28.57 (2.10$^{-3}$) | 150.00 (3.10$^{-4}$) | 0.00 (1.10$^{1}$) | 26.67 (4.10$^{-4}$) | 16.67 (2.10$^{-2}$) |
| GO:0001969 | Regulation of activation of membrane attack complex | 0.00 (1.10$^{1}$) | 28.57 (2.10$^{-3}$) | 150.00 (3.10$^{-4}$) | 0.00 (1.10$^{1}$) | 26.67 (4.10$^{-4}$) | 16.67 (2.10$^{-2}$) |
| GO:0002438 | Acute inflammatory response to antigenic stimulus | 0.00 (1.10$^{1}$) | 11.63 (9.10$^{-3}$) | 42.86 (5.10$^{-3}$) | 0.00 (1.10$^{1}$) | 10.64 (5.10$^{-3}$) | 7.02 (7.10$^{-2}$) |
| GO:0002673 | Regulation of acute inflammatory response | 0.00 (1.10$^{1}$) | 3.13 (3.10$^{-1}$) | 23.68 (1.10$^{-7}$) | 0.00 (1.10$^{1}$) | 6.12 (4.10$^{-6}$) | 5.07 (3.10$^{-5}$) |
| GO:0002866 | Positive regulation of acute inflammatory response to antigenic stimulus | 0.00 (1.10$^{1}$) | 16.67 (1.10$^{-2}$) | 75.00 (1.10$^{-3}$) | 0.00 (1.10$^{1}$) | 14.81 (5.10$^{-3}$) | 9.38 (9.10$^{-2}$) |
| GO:0002920 | Regulation of humoral immune response | 1.61 (1.10$^{1}$) | 5.20 (10.10$^{-3}$) | 31.03 (2.10$^{-8}$) | 0.00 (1.10$^{1}$) | 10.05 (4.10$^{-11}$) | 5.70 (4.10$^{-5}$) |
| GO:0002922 | Positive regulation of humoral immune response | 0.00 (1.10$^{1}$) | 12.20 (7.10$^{-3}$) | 42.86 (4.10$^{-3}$) | 0.00 (1.10$^{1}$) | 15.91 (2.10$^{-5}$) | 5.66 (3.10$^{-1}$) |
| GO:0006956 | Complement activation | 0.00 (1.10$^{1}$) | 4.40 (5.10$^{-3}$) | 28.26 (7.10$^{-12}$) | 0.00 (1.10$^{1}$) | 8.05 (4.10$^{-12}$) | 6.11 (3.10$^{-9}$) |
| GO:0019731 | Antibacterial humoral response | 29.41 (3.10$^{-3}$) | 10.20 (1.10$^{-2}$) | 0.00 (1.10$^{1}$) | 0.00 (1.10$^{1}$) | 13.21 (6.10$^{-5}$) | 4.69 (4.10$^{-1}$) |
| GO:0030449 | Regulation of complement activation | 0.00 (1.10$^{1}$) | 5.19 (4.10$^{-2}$) | 39.13 (2.10$^{-9}$) | 0.00 (1.10$^{1}$) | 10.14 (6.10$^{-9}$) | 7.30 (3.10$^{-6}$) |
| GO:0031714 | C5a anaphylatoxin chemotactic receptor binding | 0.00 (1.10$^{1}$) | 25.00 (3.10$^{-3}$) | 133.33 (3.10$^{-6}$) | 0.00 (1.10$^{1}$) | 27.78 (6.10$^{-5}$) | 18.18 (1.10$^{-3}$) |
| GO:0097278 | Complement-dependent cytotoxicity | 0.00 (1.10$^{1}$) | 16.67 (3.10$^{-3}$) | 80.00 (7.10$^{-5}$) | 0.00 (1.10$^{1}$) | 18.18 (4.10$^{-5}$) | 12.82 (2.10$^{-3}$) |
| GO:2000259 | Positive regulation of protein activation cascade | 0.00 (1.10$^{1}$) | 21.05 (5.10$^{-3}$) | 100.00 (6.10$^{-4}$) | 0.00 (1.10$^{1}$) | 19.05 (2.10$^{-3}$) | 12.00 (5.10$^{-2}$) |
| GO:0006957 | Complement activation, alternative pathway | 0.00 (1.10$^{1}$) | 7.22 (8.10$^{-3}$) | 37.50 (8.10$^{-6}$) | 0.00 (1.10$^{1}$) | 10.38 (1.10$^{-6}$) | 7.81 (4.10$^{-5}$) |
| GO:0006959 | Humoral immune response | 3.09 (1.10$^{1}$) | 3.96 (2.10$^{-3}$) | 16.88 (2.10$^{-9}$) | 0.00 (1.10$^{1}$) | 6.85 (3.10$^{-15}$) | 4.34 (8.10$^{-8}$) |
| GO:1990962 | Drug transport across blood-brain barrier | 0.00 (1.10$^{1}$) | 0.00 (1.10$^{1}$) | 0.00 (1.10$^{1}$) | 0.00 (1.10$^{1}$) | 5.56 (1.10$^{1}$) | 19.05 (2.10$^{-3}$) |
| GO:0008559 | Xenobiotic transmembrane transporting ATP | 0.00 (1.10$^{1}$) | 3.03 (1.10$^{1}$) | 0.00 (1.10$^{1}$) | 0.00 (1.10$^{1}$) | 5.56 (6.10$^{-1}$) | 11.36 (2.10$^{-3}$) |
| GO:0010866 | Regulation of triglyceride biosynthetic process | 0.00 (1.10$^{1}$) | 13.04 (2.10$^{-3}$) | 50.00 (3.10$^{-4}$) | 0.00 (1.10$^{1}$) | 16.00 (3.10$^{-6}$) | 9.84 (1.10$^{-3}$) |
| GO:0019432 | Triglyceride biosynthetic process | 2.44 (1.10$^{1}$) | 7.08 (5.10$^{-3}$) | 26.32 (4.10$^{-4}$) | 0.00 (1.10$^{1}$) | 8.87 (5.10$^{-6}$) | 7.33 (3.10$^{-5}$) |
| GO:0010884 | Positive regulation of lipid storage | 0.00 (1.10$^{1}$) | 10.53 (5.10$^{-3}$) | 40.00 (5.10$^{-4}$) | 0.00 (1.10$^{1}$) | 11.29 (2.10$^{-4}$) | 9.33 (5.10$^{-4}$) |

| | | | | | | | |
|---|---|---|---|---|---|---|---|
| GO:0016125 | Sterol metabolic process | 1.96 (1.10$^{1}$) | 3.72 (3.10$^{-3}$) | 2.74 (1.10$^{1}$) | 0.00 (1.10$^{1}$) | 4.26 (9.10$^{-6}$) | 5.64 (8.10$^{-13}$) |
| GO:0006694 | Steroid biosynthetic process | 1.91 (1.10$^{1}$) | 3.18 (2.10$^{-2}$) | 2.70 (1.10$^{1}$) | 0.00 (1.10$^{1}$) | 4.56 (9.10$^{-7}$) | 5.85 (4.10$^{-14}$) |
| GO:0043651 | Linoleic acid metabolic process | 13.33 (1.10$^{1}$) | 4.65 (9.10$^{-1}$) | 0.00 (1.10$^{1}$) | 0.00 (1.10$^{1}$) | 4.26 (8.10$^{-1}$) | 10.53 (10.10$^{-4}$) |
| GO:0042760 | Very long-chain fatty acid catabolic process | 11.11 (1.10$^{1}$) | 8.33 (5.10$^{-1}$) | 0.00 (1.10$^{1}$) | 0.00 (1.10$^{1}$) | 11.11 (7.10$^{-2}$) | 12.50 (9.10$^{-3}$) |
| GO:0033540 | Fatty acid beta-oxidation using acyl-CoA oxidase | 11.11 (1.10$^{1}$) | 4.17 (1.10$^{1}$) | 0.00 (1.10$^{1}$) | 0.00 (1.10$^{1}$) | 3.70 (1.10$^{1}$) | 15.63 (6.10$^{-4}$) |
| GO:0006699 | Bile acid biosynthetic process | 0.00 (1.10$^{1}$) | 2.25 (1.10$^{1}$) | 0.00 (1.10$^{1}$) | 0.00 (1.10$^{1}$) | 8.16 (4.10$^{-4}$) | 11.02 (1.10$^{-8}$) |
| GO:0015722 | Canalicular bile acid transport | 0.00 (1.10$^{1}$) | 0.00 (1.10$^{1}$) | 0.00 (1.10$^{1}$) | 0.00 (1.10$^{1}$) | 13.33 (8.10$^{-3}$) | 13.89 (10.10$^{-4}$) |
| GO:0034040 | Lipid-transporting ATPase activity | 11.11 (10.10$^{-1}$) | 0.00 (1.10$^{1}$) | 0.00 (1.10$^{1}$) | 0.00 (1.10$^{1}$) | 14.81 (6.10$^{-3}$) | 9.09 (6.10$^{-2}$) |
| GO:0034374 | Low-density lipoprotein particle remodeling | 0.00 (1.10$^{1}$) | 9.09 (5.10$^{-1}$) | 25.00 (10.10$^{-1}$) | 0.00 (1.10$^{1}$) | 16.67 (3.10$^{-3}$) | 10.34 (7.10$^{-2}$) |
| GO:0034379 | Very-low-density lipoprotein particle assembly | 0.00 (1.10$^{1}$) | 4.55 (1.10$^{1}$) | 0.00 (1.10$^{1}$) | 0.00 (1.10$^{1}$) | 8.33 (4.10$^{-1}$) | 17.24 (3.10$^{-4}$) |
| GO:0043020 | NADPH oxidase complex | 28.57 (10.10$^{-3}$) | 13.16 (3.10$^{-3}$) | 28.57 (5.10$^{-1}$) | 25.00 (1.10$^{1}$) | 16.67 (9.10$^{-6}$) | 11.54 (3.10$^{-4}$) |
| GO:0050664 | Oxidoreductase activity, acting on NAD(P)H oxygen as receptor | 25.00 (2.10$^{-1}$) | 18.18 (7.10$^{-3}$) | 25.00 (9.10$^{-1}$) | 50.00 (9.10$^{-1}$) | 20.83 (2.10$^{-4}$) | 16.67 (2.10$^{-4}$) |
| GO:0072376 | Protein activation cascade | 0.00 (1.10$^{1}$) | 3.93 (10.10$^{-3}$) | 25.00 (2.10$^{-11}$) | 0.00 (1.10$^{1}$) | 7.19 (4.10$^{-11}$) | 5.71 (4.10$^{-9}$) |
| GO:1903317 | Regulation of protein maturation | 0.00 (1.10$^{1}$) | 3.44 (9.10$^{-2}$) | 22.73 (2.10$^{-8}$) | 3.85 (1.10$^{1}$) | 5.57 (5.10$^{-6}$) | 4.62 (4.10$^{-5}$) |
| GO:0035375 | Zymogen binding | 60.00 (7.10$^{-3}$) | 21.43 (2.10$^{-2}$) | 0.00 (1.10$^{1}$) | 0.00 (1.10$^{1}$) | 20.00 (2.10$^{-2}$) | 11.11 (2.10$^{-1}$) |
| GO:0042588 | Zymogen granule | 16.67 (2.10$^{-2}$) | 8.82 (5.10$^{-3}$) | 0.00 (1.10$^{1}$) | 0.00 (1.10$^{1}$) | 10.53 (5.10$^{-5}$) | 8.60 (1.10$^{-4}$) |
| GO:0042589 | Zymogen granule membrane | 21.05 (2.10$^{-2}$) | 7.69 (9.10$^{-2}$) | 0.00 (1.10$^{1}$) | 0.00 (1.10$^{1}$) | 10.53 (1.10$^{-3}$) | 8.45 (2.10$^{-3}$) |
| GO:0005212 | Structural constituent of eye lens | 0.00 (1.10$^{1}$) | 2.80 (10.10$^{-1}$) | 0.00 (1.10$^{1}$) | 0.00 (1.10$^{1}$) | 11.11 (10.10$^{-8}$) | 10.42 (2.10$^{-9}$) |
| GO:0150064 | Vertebrate eye-specific patterning | 0.00 (1.10$^{1}$) | 18.18 (7.10$^{-3}$) | 75.00 (9.10$^{-4}$) | 0.00 (1.10$^{1}$) | 16.67 (3.10$^{-3}$) | 10.34 (7.10$^{-2}$) |
| GO:0098883 | Synapse pruning | 0.00 (1.10$^{1}$) | 12.50 (3.10$^{-2}$) | 60.00 (3.10$^{-3}$) | 0.00 (1.10$^{1}$) | 11.43 (2.10$^{-2}$) | 6.98 (2.10$^{-1}$) |
| GO:0150062 | Complement-mediated synapse pruning | 0.00 (1.10$^{1}$) | 18.18 (7.10$^{-3}$) | 75.00 (9.10$^{-4}$) | 0.00 (1.10$^{1}$) | 16.67 (3.10$^{-3}$) | 10.34 (7.10$^{-2}$) |
| GO:0004089 | Carbonate dehydratase activity | 25.00 (2.10$^{-3}$) | 9.09 (2.10$^{-2}$) | 0.00 (1.10$^{1}$) | 0.00 (1.10$^{1}$) | 6.67 (9.10$^{-2}$) | 5.41 (8.10$^{-2}$) |
| GO:0004866 | Endopeptidase inhibitor activity | 1.14 (1.10$^{1}$) | 3.73 (2.10$^{-3}$) | 13.64 (1.10$^{-7}$) | 4.08 (1.10$^{1}$) | 3.23 (2.10$^{-3}$) | 3.85 (10.10$^{-7}$) |
| GO:0017042 | Glycosylceramidase activity | 0.00 (1.10$^{1}$) | 14.29 (2.10$^{-1}$) | 0.00 (1.10$^{1}$) | 0.00 (1.10$^{1}$) | 26.67 (7.10$^{-4}$) | 22.22 (5.10$^{-4}$) |
| GO:0018879 | Biphenyl metabolic process | 0.00 (1.10$^{1}$) | 0.00 (1.10$^{1}$) | 0.00 (1.10$^{1}$) | 0.00 (1.10$^{1}$) | 8.33 (4.10$^{-1}$) | 13.79 (6.10$^{-3}$) |
| GO:0045745 | Positive regulation of G protein-coupled receptor signaling pathway | 0.00 (1.10$^{1}$) | 5.48 (3.10$^{-1}$) | 33.33 (1.10$^{-3}$) | 0.00 (1.10$^{1}$) | 6.25 (4.10$^{-2}$) | 4.17 (3.10$^{-1}$) |
| GO:0046581 | Intercellular canaliculus | 0.00 (1.10$^{1}$) | 2.17 (1.10$^{1}$) | 0.00 (1.10$^{1}$) | 0.00 (1.10$^{1}$) | 9.80 (5.10$^{-3}$) | 11.11 (7.10$^{-5}$) |
| GO:0052695 | Cellular glucuronidation | 0.00 (1.10$^{1}$) | 0.00 (1.10$^{1}$) | 0.00 (1.10$^{1}$) | 50.00 (1.10$^{1}$) | 4.17 (1.10$^{1}$) | 13.79 (6.10$^{-3}$) |
| GO:0071276 | Cellular response to cadmium ion | 0.00 (1.10$^{1}$) | 4.88 (8.10$^{-1}$) | 42.86 (4.10$^{-3}$) | 25.00 (1.10$^{1}$) | 4.55 (7.10$^{-1}$) | 1.89 (1.10$^{1}$) |
| GO:1905153 | Regulation of membrane invagination | 0.00 (1.10$^{1}$) | 10.53 (4.10$^{-2}$) | 50.00 (4.10$^{-3}$) | 0.00 (1.10$^{1}$) | 9.76 (3.10$^{-2}$) | 6.00 (2.10$^{-1}$) |
| GO:1990578 | Perinuclear endoplasmic reticulum membrane | 14.29 (1.10$^{1}$) | 10.53 (4.10$^{-1}$) | 33.33 (1.10$^{1}$) | 0.00 (1.10$^{1}$) | 14.29 (3.10$^{-2}$) | 15.38 (2.10$^{-3}$) |
| GO:2000425 | Regulation of apoptotic cell clearance | 0.00 (1.10$^{1}$) | 18.18 (7.10$^{-3}$) | 75.00 (9.10$^{-4}$) | 0.00 (1.10$^{1}$) | 16.67 (3.10$^{-3}$) | 10.34 (7.10$^{-2}$) |

**S2 Table:** The top 10 most overrepresented GO terms from the differentially expressed gene sets (FDR <0.05 and |LFC| >1), according to light treatment comparisons and timepoints.

| | | **GO** | **Annotation** | **Odds-ratio (FDR)** | **Differentially expressed genes (downregulated under higher light level)** |
|---|---|---|---|---|---|
| **At night-time** | **0.1 lux *vs* control** | GO:0035375 | Zymogen binding | 60.00 ($7.10^{-3}$) | 3 (3) |
| | | GO:0019731 | Antibacterial humoral response | 29.41 ($3.10^{-3}$) | 5 (5) |
| | | GO:0043020 | NADPH oxidase complex | 28.57 ($10.10^{-3}$) | 4 (2) |
| | | GO:0004089 | Carbonate dehydratase activity | 25.00 ($2.10^{-3}$) | 5 (5) |
| | | GO:0017171 | Serine hydrolase activity | 4.78 ($9.10^{-3}$) | 11 (9) |
| | | GO:0006952 | Defence response | 2.74 ($1.10^{-3}$) | 32 (23) |
| | **5 lux *vs* control** | GO:0001969 | Regulation of activation of membrane attack complex | 28.57 ($2.10^{-3}$) | 4 (4) |
| | | GO:0031714 | C5a anaphylatoxin chemotactic receptor binding | 25.00 ($3.10^{-3}$) | 4 (4) |
| | | GO:2000425 | Regulation of apoptotic cell clearance | 18.18 ($7.10^{-3}$) | 4 (4) |
| | | GO:0050664 | Oxidoreductase activity, acting on NAD(P)H, oxygen as receptor | 18.18 ($7.10^{-3}$) | 4 (3) |
| | | GO:0097278 | Complement-dependent cytotoxicity | 16.67 ($3.10^{-3}$) | 5 (5) |
| | | GO:0043020 | NADPH oxidase complex | 13.16 ($2.10^{-3}$) | 5 (4) |
| | | GO:0010866 | Regulation of triglyceride biosynthetic process | 13.04 ($2.10^{-3}$) | 6 (6) |
| | | GO:0010884 | Positive regulation of lipid storage | 10.53 ($5.10^{-3}$) | 6 (6) |
| | | GO:0042588 | Zymogen granule | 8.82 ($5.10^{-3}$) | 6 (6) |
| | | GO:0030049 | Muscle filament sliding | 6.30 ($7.10^{-3}$) | 8 (1) |
| | **5 lux *vs* 0.1 lux** | GO:0031714 | C5a anaphylatoxin chemotactic receptor binding | 133.33 ($3.10^{-6}$) | 4 (4) |
| | | GO:0097278 | Complement-dependent cytotoxicity | 80.00 ($7.10^{-5}$) | 4 (4) |
| | | GO:2000425 | Regulation of apoptotic cell clearance | 75.00 ($9.10^{-4}$) | 3 (3) |
| | | GO:0098883 | Synapse pruning | 60.00 ($3.10^{-3}$) | 3 (3) |
| | | GO:1905153 | Regulation of membrane invagination | 50.00 ($4.10^{-3}$) | 3 (3) |
| | | GO:0071276 | Cellular response to cadmium ion | 42.86 ($4.10^{-3}$) | 3 (3) |
| | | GO:0010884 | Positive regulation of lipid storage | 40.00 ($5.10^{-4}$) | 4 (4) |
| | | GO:0045745 | Positive regulation of G protein-coupled receptor signalling pathway | 33.33 ($1.10^{-3}$) | 4 (4) |
| | | GO:0006956 | Complement activation | 28.26 ($7.10^{-12}$) | 13 (12) |
| | | GO:0019432 | Triglyceride biosynthetic process | 26.32 ($4.10^{-4}$) | 5 (5) |
| **At daytime** | **5 lux *vs* control** | GO:0031714 | C5a anaphylatoxin chemotactic receptor binding | 27.78 ($6.10^{-5}$) | 6(5) |
| | | GO:0001798 | Positive regulation of type IIa hypersensitivity | 26.67 ($4.10^{-4}$) | 5(4) |
| | | GO:0001905 | Activation of membrane attack complex | 26.67 ($4.10^{-4}$) | 5(4) |
| | | GO:0017042 | Glycosylceramidase activity | 26.67 ($7.10^{-4}$) | 5(4) |
| | | GO:0050664 | Oxidoreductase activity, acting on NAD(P)H, oxygen as acceptor | 20.83 ($2.10^{-4}$) | 8(5) |
| | | GO:2000259 | Positive regulation of protein activation cascade | 19.05 ($2.10^{-3}$) | 7(4) |
| | | GO:0097278 | Complement-dependent cytotoxicity | 18.18 ($4.10^{-5}$) | 11(6) |
| | | GO:0002524 | Hypersensitivity | 16.67 ($3.10^{-3}$) | 8(4) |
| | | GO:0034374 | Low-density lipoprotein particle remodelling | 16.67 ($3.10^{-3}$) | 8(4) |
| | | GO:0150062 | Complement-mediated synapse pruning | 16.67 ($3.10^{-3}$) | 8(4) |
| | **5 lux *vs* 0.1 lux** | GO:0017042 | Glycosylceramidase activity | 22.22 ($5.10^{-4}$) | 5(4) |
| | | GO:1990962 | Xenobiotic transport across blood-brain barrier | 19.05 ($2.10^{-3}$) | 6(4) |
| | | GO:0031714 | C5a anaphylatoxin chemotactic receptor binding | 18.18 ($1.10^{-3}$) | 6(4) |
| | | GO:0034379 | Very-low-density lipoprotein particle assembly | 17.24 ($3.10^{-4}$) | 8(5) |
| | | GO:0050664 | Oxidoreductase activity, acting on NAD(P)H, oxygen as acceptor | 16.67 ($2.10^{-4}$) | 8(5) |
| | | GO:0033540 | Fatty acid beta-oxidation using acyl-CoA oxidase | 15.63 ($6.10^{-4}$) | 9(5) |
| | | GO:1990578 | Perinuclear endoplasmic reticulum membrane | 15.38 ($2.10^{-3}$) | 7(4) |
| | | GO:0015722 | Canalicular bile acid transport | 13.89 ($10.10^{-3}$) | 10(5) |
| | | GO:0018879 | Biphenyl metabolic process | 13.79 ($6.10^{-3}$) | 8(4) |
| | | GO:0052695 | Cellular glucuronidation | 13.79 ($6.10^{-3}$) | 8(4) |

**S3 Table:** Enrichment level of GO terms related to circadian rhythm according to light treatment comparisons and timepoints.

| | | **At night-time** | | | **At daytime** | | |
|---|---|---|---|---|---|---|---|
| | | **0.1 lux *vs* control** | **5 lux *vs* control** | **5 lux *vs* 0.1 lux** | **0.1 lux *vs* control** | **5 lux *vs* control** | **5 lux *vs* 0.1 lux** |
| **GO** | **Annotation** | **Odds-ratio (FDR)** | **Odds-ratio (FDR)** | **Odds-ratio (FDR)** | **Odds-ratio (FDR)** | **Odds-ratio (FDR)** | **Odds-ratio (FDR)** |
| GO:0007622 | Rhythmic behavior | 2.78 ($1.10^{1}$) | 1.00 ($1.10^{1}$) | 0.00 ($1.10^{1}$) | 0.00 ($1.10^{1}$) | 0.00 ($1.10^{1}$) | 0.00 ($1.10^{1}$) |
| GO:0007623 | Circadian rhythm | 0.57 ($1.10^{1}$) | 0.40 ($1.10^{1}$) | 1.19 ($1.10^{1}$) | 0.00 ($1.10^{1}$) | 0.18 ($1.10^{1}$) | 0.77 ($1.10^{1}$) |
| GO:0009416 | Response to light stimulus | 0.00 ($1.10^{1}$) | 0.30 ($1.10^{1}$) | 0.88 ($1.10^{1}$) | 2.99 ($1.10^{1}$) | 0.95 ($1.10^{1}$) | 0.79 ($1.10^{1}$) |
| GO:0009649 | Entrainment of circadian clock | 0.00 ($1.10^{1}$) | 0.00 ($1.10^{1}$) | 0.00 ($1.10^{1}$) | 0.00 ($1.10^{1}$) | 0.00 ($1.10^{1}$) | 0.00 ($1.10^{1}$) |
| GO:0010840 | Regulation of circadian sleep/wake cycle, wakefulness | 0.00 ($1.10^{1}$) | 0.00 ($1.10^{1}$) | 0.00 ($1.10^{1}$) | 0.00 ($1.10^{1}$) | 0.00 ($1.10^{1}$) | 0.00 ($1.10^{1}$) |
| GO:0022410 | Circadian sleep/wake cycle process | 0.00 ($1.10^{1}$) | 0.00 ($1.10^{1}$) | 0.00 ($1.10^{1}$) | 0.00 ($1.10^{1}$) | 0.00 ($1.10^{1}$) | 0.00 ($1.10^{1}$) |
| GO:0032922 | Circadian regulation of gene expression | 0.00 ($1.10^{1}$) | 0.00 ($1.10^{1}$) | 3.57 ($1.10^{1}$) | 0.00 ($1.10^{1}$) | 0.55 ($1.10^{1}$) | 0.45 ($1.10^{1}$) |
| GO:0042320 | Regulation of circadian sleep/wake cycle, REM sleep | 0.00 ($1.10^{1}$) | 0.00 ($1.10^{1}$) | 0.00 ($1.10^{1}$) | 0.00 ($1.10^{1}$) | 0.00 ($1.10^{1}$) | 0.00 ($1.10^{1}$) |
| GO:0042745 | Circadian sleep/wake cycle | 0.00 ($1.10^{1}$) | 0.00 ($1.10^{1}$) | 0.00 ($1.10^{1}$) | 0.00 ($1.10^{1}$) | 0.00 ($1.10^{1}$) | 0.00 ($1.10^{1}$) |
| GO:0042746 | Circadian sleep/wake cycle, wakefulness | 0.00 ($1.10^{1}$) | 0.00 ($1.10^{1}$) | 0.00 ($1.10^{1}$) | 0.00 ($1.10^{1}$) | 0.00 ($1.10^{1}$) | 0.00 ($1.10^{1}$) |
| GO:0042747 | Circadian sleep/wake cycle, REM sleep | 0.00 ($1.10^{1}$) | 0.00 ($1.10^{1}$) | 0.00 ($1.10^{1}$) | 0.00 ($1.10^{1}$) | 0.00 ($1.10^{1}$) | 0.00 ($1.10^{1}$) |
| GO:0042749 | Regulation of circadian sleep/wake cycle | 0.00 ($1.10^{1}$) | 0.00 ($1.10^{1}$) | 0.00 ($1.10^{1}$) | 0.00 ($1.10^{1}$) | 0.00 ($1.10^{1}$) | 0.00 ($1.10^{1}$) |
| GO:0042752 | Regulation of circadian rhythm | 1.16 ($1.10^{1}$) | 0.83 ($1.10^{1}$) | 0.00 ($1.10^{1}$) | 0.00 ($1.10^{1}$) | 0.00 ($1.10^{1}$) | 0.63 ($1.10^{1}$) |
| GO:0042753 | Positive regulation of circadian rhythm | 0.00 ($1.10^{1}$) | 0.00 ($1.10^{1}$) | 0.00 ($1.10^{1}$) | 0.00 ($1.10^{1}$) | 0.00 ($1.10^{1}$) | 0.00 ($1.10^{1}$) |
| GO:0042754 | Negative regulation of circadian rhythm | 0.00 ($1.10^{1}$) | 0.00 ($1.10^{1}$) | 0.00 ($1.10^{1}$) | 0.00 ($1.10^{1}$) | 0.00 ($1.10^{1}$) | 0.00 ($1.10^{1}$) |
| GO:0043153 | Entrainment of circadian clock by photoperiod | 0.00 ($1.10^{1}$) | 0.00 ($1.10^{1}$) | 0.00 ($1.10^{1}$) | 0.00 ($1.10^{1}$) | 0.00 ($1.10^{1}$) | 0.00 ($1.10^{1}$) |
| GO:0045187 | Regulation of circadian sleep/wake cycle, sleep | 0.00 ($1.10^{1}$) | 0.00 ($1.10^{1}$) | 0.00 ($1.10^{1}$) | 0.00 ($1.10^{1}$) | 0.00 ($1.10^{1}$) | 0.00 ($1.10^{1}$) |
| GO:0045475 | Locomotor rhythm | 0.00 ($1.10^{1}$) | 0.00 ($1.10^{1}$) | 0.00 ($1.10^{1}$) | 0.00 ($1.10^{1}$) | 0.00 ($1.10^{1}$) | 0.00 ($1.10^{1}$) |
| GO:0045938 | Positive regulation of circadian sleep/wake cycle | 0.00 ($1.10^{1}$) | 0.00 ($1.10^{1}$) | 0.00 ($1.10^{1}$) | 0.00 ($1.10^{1}$) | 0.00 ($1.10^{1}$) | 0.00 ($1.10^{1}$) |
| GO:0048511 | Rhythmic process | 0.40 ($1.10^{1}$) | 0.43 ($1.10^{1}$) | 0.85 ($1.10^{1}$) | 0.00 ($1.10^{1}$) | 0.13 ($1.10^{1}$) | 0.76 ($1.10^{1}$) |
| GO:0048512 | Circadian behavior | 2.78 ($1.10^{1}$) | 1.00 ($1.10^{1}$) | 0.00 ($1.10^{1}$) | 0.00 ($1.10^{1}$) | 0.00 ($1.10^{1}$) | 0.00 ($1.10^{1}$) |
| GO:0050802 | Circadian sleep/wake cycle, sleep | 0.00 ($1.10^{1}$) | 0.00 ($1.10^{1}$) | 0.00 ($1.10^{1}$) | 0.00 ($1.10^{1}$) | 0.00 ($1.10^{1}$) | 0.00 ($1.10^{1}$) |
| GO:0060024 | Rhythmic synaptic transmission | 0.00 ($1.10^{1}$) | 0.00 ($1.10^{1}$) | 0.00 ($1.10^{1}$) | 0.00 ($1.10^{1}$) | 0.00 ($1.10^{1}$) | 0.00 ($1.10^{1}$) |
| GO:1904059 | Regulation of locomotor rhythm | 0.00 ($1.10^{1}$) | 0.00 ($1.10^{1}$) | 0.00 ($1.10^{1}$) | 0.00 ($1.10^{1}$) | 0.00 ($1.10^{1}$) | 0.00 ($1.10^{1}$) |

**S1 Appendix: Sequencing and assembly of *B. bufo de novo* transcriptome**

To maximize the set of described genes and to provide a guide for transcriptome assembly, we built a transcriptome assembly from our samples as well as from nine *B. bufo* tadpole brains, sampled at various development stages (tadpoles and juveniles). Total RNA from the latter samples was extracted by adding TRI Reagent (Molecular Research Center MRC, TR118) to the samples and by homogenizing tissues with a piston. All remaining steps were carried out according to the manufacturers' protocols (Molecular Research Center MRC, TR118). Then, the RNA was treated with Turbo DNase enzyme (Turbo DNA free kit, Invitrogen, AM1907) and assayed by fluorescence using Qubit fluorometer (Qubit® RNA HS Assay Kits Molecular Probes, Invitrogen, Q32855). The quality was checked by Bioanalyzer 2100 with Agilent RNA 6000 Nano Kit (Agilent, 5067-1511). mRNA was isolated and pooled to allow the construction of one cDNA library using TruSeq™ RNA Sample Prep Kit v2 -Set A (ILLUMINA, RS-122-2001). cDNA concentration quality was checked using the previous Bioanalyzer 2100 with High Sensitivity DNA Assay kit (Agilent, 5067-4626). Library was sequenced with an Illumina® HiSeq4000™ machine on the GenomEast platform hosted at the IGBMC (GenomEast Platform – IGBMC, Illkirch, France), resulting in paired-end reads with reads 100 bp long.
